# The relation between intrinsic protein conformational changes and ligand binding

**DOI:** 10.1101/2020.03.03.974634

**Authors:** Marijn de Boer

**Affiliations:** Molecular Microscopy Research Group, Zernike Institute for Advanced Material, University of Groningen, 9747 AG Groningen, The Netherlands

## Abstract

Structural changes in proteins allow them to exist in several conformations. Non-covalent interactions with ligands drive the structural changes, thereby allowing the protein to perform its biological function. Recent findings suggest that many proteins are always in an equilibrium of different conformations and that each of these conformations can be formed by both the ligand-free and ligand-bound protein. By using classical statistical mechanics, we derived the equilibrium probabilities of forming a conformation with and without ligand. We found, under certain conditions, that increasing the probability of forming a conformation by the ligand-free protein also increases the probability of forming the same conformation when the protein has a ligand bound. Further, we found that changes in the conformational equilibrium of the ligand-free protein can increase or decrease the affinity for the ligand.

## 2 INTRODUCTION

The polypeptide of proteins-except intrinsically disordered proteins-folds into a three-dimensional structure. This structure is not rigid but can change in response to specific interactions with ligands^1^. For many proteins these structural changes involve the collective motion of many atoms, such as the complete reorientation of a domain. In others, more localized changes occur, leading to changes in the relative position of only a few residues, such as a loop displacement.

The coupling of ligand binding to protein conformational changes is fundamental to almost all biological processes. For instance, signalling proteins switch from an inactive to an active conformation upon the binding of an effector molecule^2^. The conformational change of the signalling protein can be used to transmit a signal to downstream members of the signalling pathway. In enzymes, the conformational changes bring the catalytic residues in the proper orientation and allow the substrate to bind and the product to be released^3^. In membrane transporter proteins (e.g., ATP-binding cassette (ABC) transporters), switching between an inward- and outward-facing conformation exposes a substrate-binding site on alternative sides of the membrane^4^, thereby providing a mechanism to translocate compounds across the membrane.

The mechanism in which ligand interactions drive the structural change is termed the induced-fit mechanism. In the classical induced-fit mechanism^5^, a ligand-free protein is in a single conformation and upon binding of the ligand or substrate, a new conformation is formed. However, nuclear magnetic resonance (NMR)^6-8^, electron paramagnetic resonance (EPR)^9^, single-molecule Förster resonance energy transfer (smFRET)^10-19^ and other data^20-22^, indicate that proteins sample a range of conformations with and without ligand bound. This led to the notion that, proteins are inherently dynamic and are always in an equilibrium of different conformations^10, 23-25^. Most interestingly, for many proteins, experimental^7, 8, 12, 19, 26-30^ and computational^31-33^ results suggest that the structure of the ligand-free and ligand-bound conformations are highly similar. Thus, in contrast to the classical induced-fit mechanism, ligand interactions do not induce new conformations in these cases, but only redistribute the conformational equilibrium that already exists in the ligand-free protein^23-25^. Examples of proteins with such an intrinsic conformational equilibrium include the substrate-binding proteins (SBPs) of ABC importers^10, 11^, and the soluble ABC protein ABCE1^12^. Other prominent examples are the proteins adenylate kinase^18^, RNase A^7^, dihydrofolate reductase^29^, ubiquitin^30^, SecA^28^, the Lac repressor^26^ and DNA polymerase^16, 17^ and many others^25^.

In this work, classical statistical mechanics is used to describe the conformational ensemble of a protein that binds a single ligand. We derived the equilibrium probabilities of forming a conformation with and without ligand. By assuming that the Hamiltonian is additive, these probabilities can be related to each other. We found that in particular cases, increasing the probability of forming a conformation without ligand also increases the probability of forming the same conformation with ligand. Moreover, the dissociation constant K_D_ between the protein and the ligand is found to be sensitive to changes in the conformational equilibrium of the ligand-free protein.

## 3 RESULTS

### 3.1 The ligand-free and ligand-bound phase space density

We begin with a description of the protein in the absence of ligand, i.e., the apo protein. In a classical system, each microscopic state can be specified by the position and momentum of the *N* atoms of the protein, denoted by the 3*N*-dimensional vectors ***x*** and ***p***, respectively. In the canonical ensemble^34^, the phase space density of the ligand-free protein is

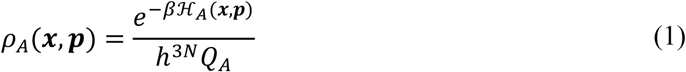

where ℋ_*A*_(***x, p***) is the ligand-free (apo) protein Hamiltonian function, *h* is the Planck constant and *β* = 1/*k*_*B*_*T*, where *k*_*B*_ is the Boltzmann constant and *T* the absolute temperature. The canonical partition function *Q*_*A*_ is

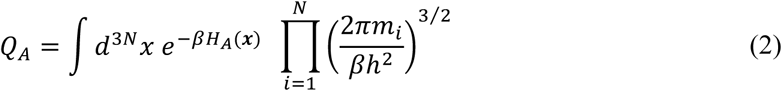

where *H*_*A*_(***x***) is the configurational part of ℋ_*A*_(***x, p***) and *m*_*i*_ is the mass of the *i*th atom. We are interested in the conformational behaviour of the protein, so we integrate out the momentum degrees of freedom to obtain the configurational space density,

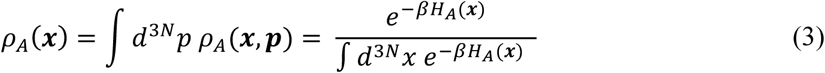

In an analogous manner as we did for the protein, we can specify the ligand molecule by the position and momentum of the *M* atoms of the ligand, denoted by the 3*M*-dimensional vectors ***x***′ and ***p***′, respectively. The phase space density of the ligand-bound protein is

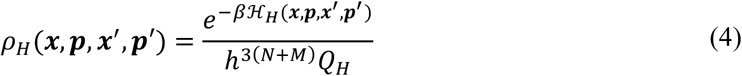

where ℋ_*H*_(***x, p, x***′, ***p***′) is the Hamiltonian of the ligand-bound (holo) protein. The canonical partition function of the ligand-bound protein *Q*_*H*_ is

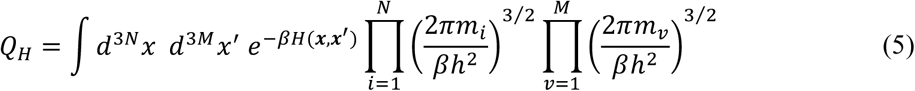

where *m*_*i*_ and *m*_*v*_ are the masses of the *i*th and *v*th atom of the protein and the ligand, respectively, and *H*_*H*_ (***x, x***′) is the configurational part of ℋ_*H*_(***x, p, x***′, ***p***′).

If the Hamiltonian function is additive, we can express *H*_*H*_(***x, x***′) as

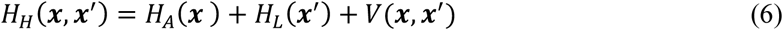

where *H*_*A*_(***x***) and *H*_*L*_(***x***′) are the configurational parts of the Hamiltonian of the ligand-free protein and the free ligand, respectively. *V*(***x, x***′) is the interaction potential between the protein and the ligand. The separation of *H*_*H*_ (***x, x***′) holds for all pairwise additive potentials, but also for non-pairwise additive potentials, such as three- or higher-order body forces^35^. For example, in the presence of three-body forces: *H*_*H*_ (***x, x***′) = (*H*_*PP*_ + *H*_*PPP*_) + (*H*_*LL*_ + *H*_*LLL*_) + (*H*_*PL*_ + *H*_*PPL*_ + *H*_*PLL*_), where *H*_*ij*_ and *H*_*ijk*_ are the pairwise and three-body interaction potentials, respectively, between the protein (P) and ligand (L) coordinates.

We obtain the configurational space density of the ligand-bound protein by integrating out the position and momentum degrees of freedom of the ligand and the momentum degrees of freedom of the protein,

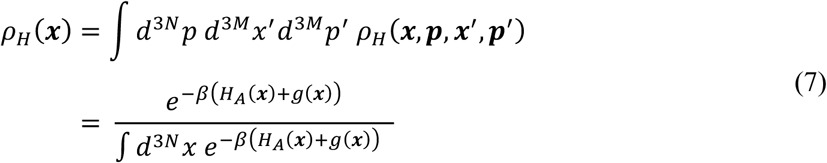

with

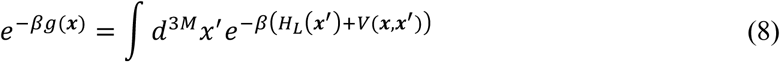

where *H*_*A*_(***x***) + *g*(***x***) is an effective Hamiltonian function that depends on the protein configurational degrees of freedom and on temperature.

The integrals of Eq. 5, 7 and 8 should be restricted to configurations that give rise to a protein-ligand complex. The definition of this complex is, to some extent, arbitrary and has been discussed in a number of studies^36-39^. For proteins that have a localized binding site, a bound complex can be defined by the configurations that have the ligand localized within a specified region of the binding site^36-39^. This region should be large enough so that it contains the most important configurations of the bound complex, i.e., those configurations with *V*(***x, x***′) ≪ 0, and should not contain a large number of configurations that are unbound, i.e., those configurations with *V*(***x, x***′) ⪆ 0^40^.

### 3.2 The protein conformation

It has been observed that many proteins exist in several conformations, that can be formed both with and without ligand present in its binding site^7-12, 19, 26-30, 41-44^(reviewed in ref^25^). Here, we derive the probabilities of forming the conformations with and without ligand, which defines the ligand-bound and ligand-free conformational equilibrium of the protein, respectively. First, we need to define a protein conformation in the context of classical statistical mechanics. Let Ω denote the set of all configurational coordinates that contribute to the ligand-free or ligand-bound partition function or to both. These configurations are often termed the native protein configurations. A protein ‘conformation’ can be defined as a confined region within the configurational space, i.e., Ω_*i*_ ⊂ Ω (Figure 1). These regions are of low free energy, with different conformations being separated by free energy barriers. In general, proteins exist in only a few conformations that are functionally and structurally distinct. The idea of course-graining the configurational space into only a few relevant degrees of freedom has been used before in other studies^45-48^. Please note that we term ***x*** a *configuration* and the set of configurations in Ω_*i*_ is termed a *conformation*.

**Figure 1.**
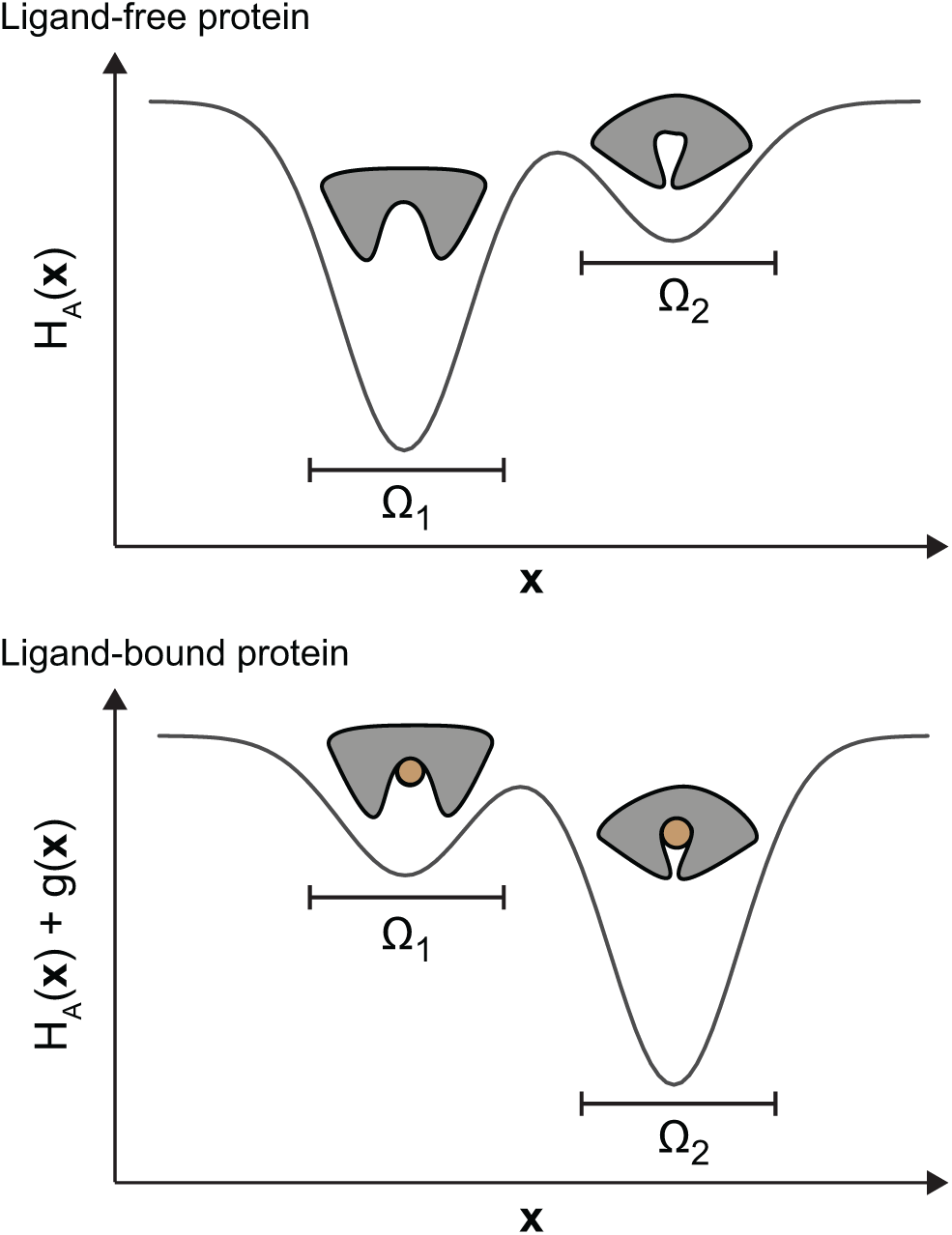
Energy landscape and the protein conformations. Schematic of the energy landscape of the ligand-free (top) and ligand-bound (bottom) protein. The protein can acquire two conformations denoted by conformation 1 and 2. The set of configurations ***x*** that belong to conformation 1 and 2 are *Ω*_1_ and *Ω*_2_, respectively. The probability to form conformation 1 with and without ligand is 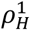 and 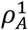, respectively, and to form conformation 2 with and without ligand is 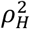 and 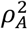, respectively. We have 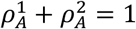 and 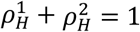.

If the native protein conformational ensemble can be described by *m* conformations (*m* ≥ 1), then the equilibrium probability that the ligand-free protein is in the *i*th conformation (*i* ∈ {1, …, *m*}) is

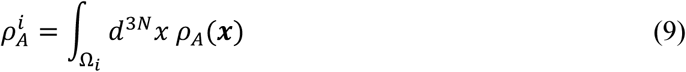

To form the *i*th conformation when the protein has a ligand bound is

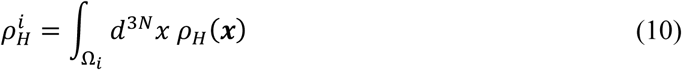

Each native protein configuration belongs to only one conformation, so for every *i* ≠ *j* we have Ω_*i*_ ⋂ Ω_*j*_ = ø and thus 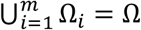, so that the probabilities are normalized to one, i.e., 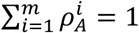 and 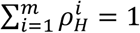.

In concluding this section, we note that in our derivation we do not assume, nor imply that the ‘measured structure’ of the *i*th conformation with and without ligand bound are identical. The measured structure of the *i*th conformation corresponds to the average configuration over the set Ω_*i*_, with the translational and rotational degrees of freedom integrated out. Integrating out the translational and rotational degrees of freedom from ***x***, gives rise to 3*N*-6 internal positional degrees of freedom ***r***. The measured structure of the *i*th conformation of the ligand-free protein then corresponds to

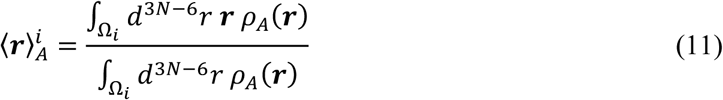

while

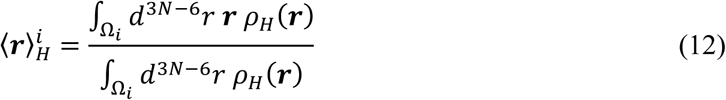

when the ligand is bound. The Jacobian for the transformation is approximately constant^35^ and therefore cancels in the division of Eq. 11 and 12. Since the phase space densities *ρ*_*A*_(***r***) and *ρ*_*H*_(***r***) can be different for ***r*** ∈ Ω_*i*_, also differences can exist between 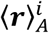 and 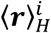. An exception is the case when *g*(***r***) (Eq. 7) varies only minimally over Ω_*i*_, than 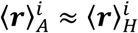. However, irrespective of 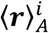 and 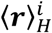, the equilibrium probability that the protein is in Ω_*i*_ with and without ligand can be calculated with Eq. 9 and 10.

### 3.3 Altering the conformational equilibrium

Next, we establish how changes in the interactions within the protein, i.e., the intra-protein interactions, affect the conformational equilibria. For that, we will analyse how 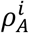 and 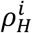 are changed when *H*_*A*_(***x***) is changed by a small perturbation *ϵθ*(***x***), with *ϵ* being a small positive number, and *θ*(***x***) is a function with units energy. The new configurational part of the Hamiltonian of the ligand-free protein is

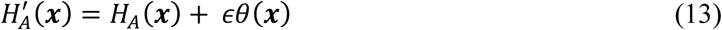

When only the intra-protein interactions are altered, while leaving the protein-ligand interactions unaltered (i.e., *H*_*L*_(***x***′) and *V*(***x, x***′) in Eq. 6 remain the same), then the new configurational part of the Hamiltonian of the ligand-bound protein is

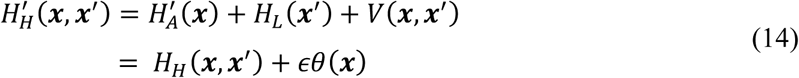

and the new effective Hamiltonian is 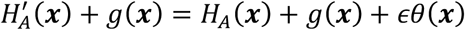.

The meaning of *ϵθ*(***x***) can be best explained trough an example (Figure 2). Suppose a protein exists in two conformations, termed conformation 1 and 2. Now, suppose that a mutation in the protein shifts the energy of each configuration of conformation 2 by a constant *ϵδ* (*ϵ* > 0), while the energy of the configurations of conformation 1 remains unaltered. In this case we have

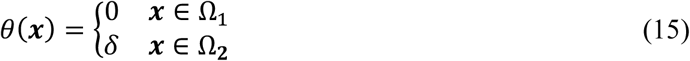

**Figure 2.**
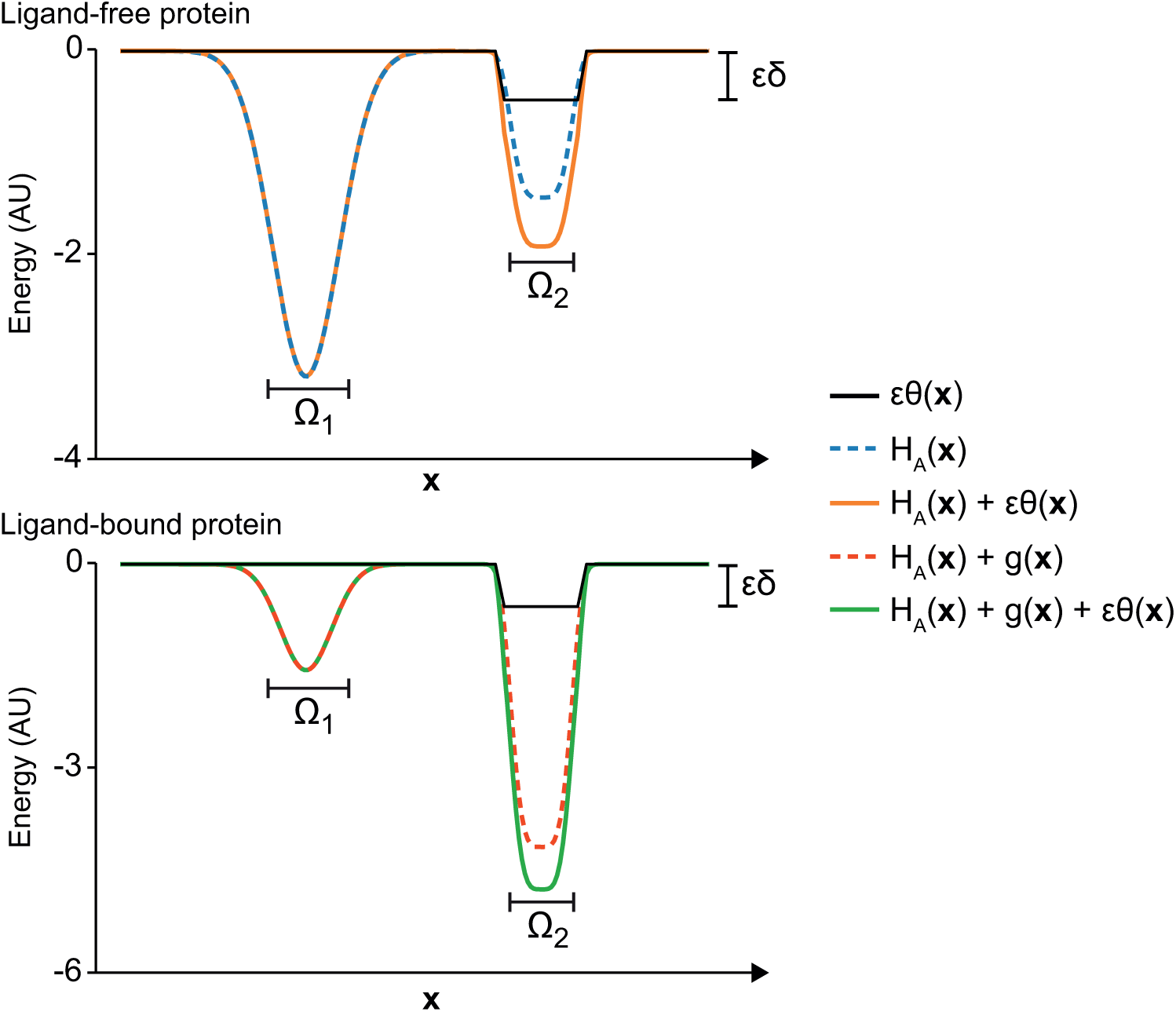
Altering the energy landscape of the ligand-free and ligand-bound protein. Example of an energy landscape of the ligand-free (top) and ligand-bound (bottom) protein as discussed in Section 3.3. The protein can acquire two conformations denoted by conformation 1 and 2. The set of configurations ***x*** that belong to conformation 1 and 2 are *Ω*_1_ and *Ω*_2_, respectively. A (hypothetical) mutation alters the energy of the configurations of conformation 2 by a constant *ϵδ* < 0. The energy landscape of the wild type ligand-free and ligand bound protein are shown in blue and red, respectively, and the mutant ligand-free and ligand bound protein are shown in orange and green, respectively.

In Figure 2 the case of *δ* < 0 is shown. The configurational part of the Hamiltonian of the mutated ligand-free protein is

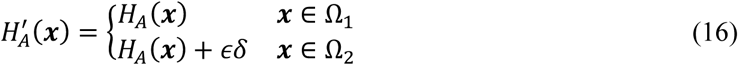

where, in this example, *H*_*A*_(***x***) is the configurational part of the Hamiltonian of the wild type ligand-free protein. If the mutation does not influence the interactions with the ligand, we have that the effective Hamiltonian of the mutated ligand-bound protein is

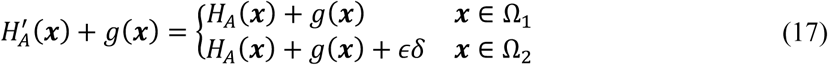

where, in this example, *H*_*A*_(***x***) + *g*(***x***) is the effective Hamiltonian of the wild type ligand-bound protein. The energy of conformation 2 will increase when *δ* > 0 and decrease when *δ* < 0. We can ask the question, how does the mutation influence the ligand-free and ligand-bound conformational equilibrium? The answer is quite simple in this example, however, it becomes more complex when *θ*(***x***) is not constant over Ω_*i*_.

In the remainder of this section we will analyse how the ligand-free and ligand-bound conformational equilibria are changed for any *θ*(***x***). First, we will establish how changes in the Hamiltonian as given by Eq. 13 affect 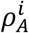. The new ligand-free conformational equilibrium becomes

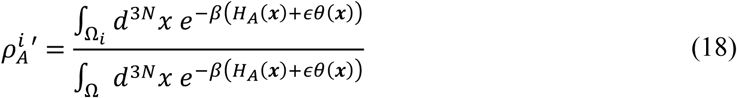

where 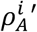 is the new equilibrium probability of forming the *i*th conformation without ligand. Expanding the exponent of Eq. 18 to first-order in *ϵ* gives

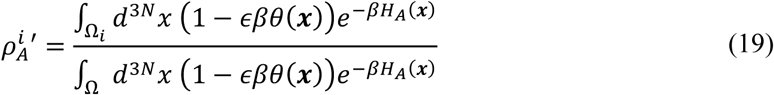

and keeping only terms up to first-order in *ϵ* results in

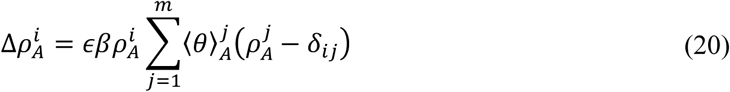

where 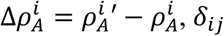 is the Kronecker delta and

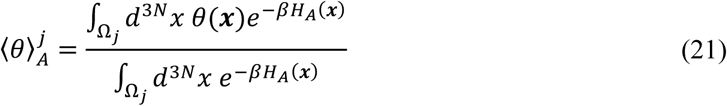

Eq. 20 shows that the new conformational equilibrium of the ligand-free protein is determined by the 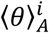 and 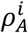 values of the *m* conformations. It can be shown that 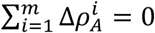, so the probabilities remain normalized when they are changed according to Eq. 20.

In a similar approach as has been done for the ligand-free conformational equilibrium, the new ligand-bound conformational equilibrium is

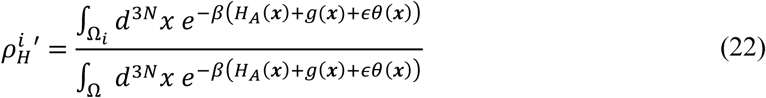

where 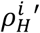 is the new equilibrium probability of forming the *i*th conformation with ligand, when the Hamiltonian is given by Eq. 14. Similar as for the ligand-free protein we obtain to first-order in *ϵ*

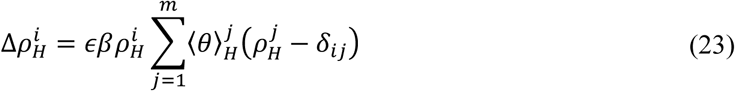

where 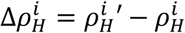 and

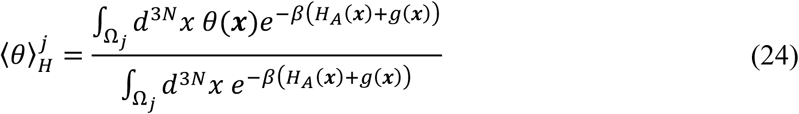

It can be shown that 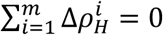. From 23 it follows that the new conformational equilibrium of the ligand-bound protein is determined by the 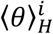 and 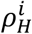 values of the *m* conformations.

In case of the example of Eq. 15, we have

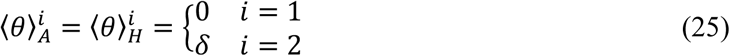

So from Eq. 20, 23 and 25 it follows that by the mutation the probabilities to form conformation *i* without and with ligand is

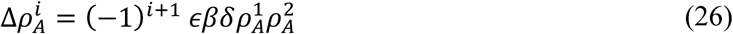

and

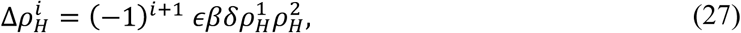

respectively. In Figure 2, the energy of conformation 2 decreases (*δ* < 0), so from Eq. 26 and 27 it follows that 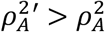 and 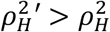. In other words, both the ligand-free and ligand-bound conformational equilibria are shifted towards conformation 2. If instead the energy of conformation 2 increases (*δ* > 0), we find that 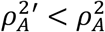 and 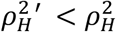, so both the ligand-free and ligand-bound conformational equilibria are shifted towards conformation 1.

### 3.4 Altering the energy of the *k*th conformation

Hereinafter, we will focus on the situation that the energy of only one conformation is altered. The conformation that is altered is denoted as the *k*th conformation, where *k* ∈ {1, …, *m*}. Formally, in this situation: *θ*(***x***) = 0 for ***x*** ∉ Ω_*k*_ and *θ*(***x***) ≠ 0 for some ***x*** ∈ Ω_*k*_. Note that *θ*(***x***) as given by Eq. 15 satisfies this criteria.

The energies of the configurations of the *k*th conformation are changed, while the energies of all the other configurations remains unchanged, when only the intra-protein interactions of the *k*th conformation are altered and all interactions that are shared with or are unique to the other conformations remain unaltered. These intra-protein interactions include those that are made when the *k*th conformation is formed and are absent in all the other conformations. Such interactions have been identified for many proteins and are most often crucial for their function and are highly conserved. For instance, residues of the MalE protein that are part of the so-called ‘balancing interface’ provide the interactions to stabilize the closed conformation of MalE and are absent in the other conformation (the open conformation)^49^.

Irrespective of the precise molecular interpretation, when only the energy of the *k*th conformation is altered, we have for *i* ≠ *k*, that

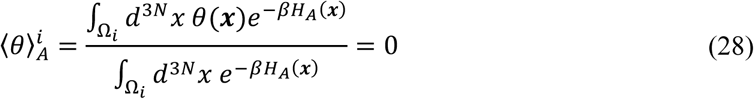

and

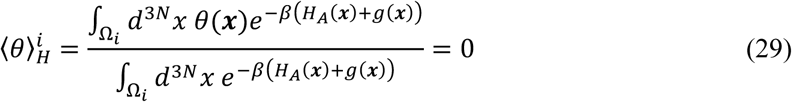

since by definition *θ*(***x***) = 0 for every ***x*** ∉ Ω_*k*_. Note that 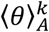 and 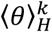 can be non-zero, since by definition *θ*(***x***) ≠ 0 for some ***x*** ∈ Ω_*k*_. From Eq. 20 and 28 it follows that

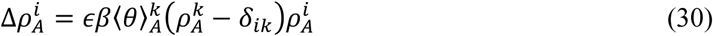

and from Eq. 23 and 29 we have

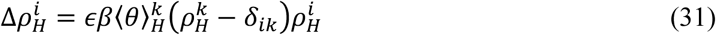

We will use Eq. 30 and 31 as a starting point to analyse how a shift in the ligand-free conformational equilibrium, which is caused by altering the energy of single conformation, will affect the ligand-bound conformational equilibrium.

### 3.5 Relating the ligand-bound and ligand-free conformational equilibrium

Now that we know how 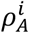 and 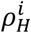 change when the intra-protein interactions of the *k*th conformation are altered (Section 3.4), we can analyse how changes in 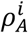 will affect 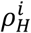. For that, we calculate the ratio 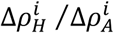. We are particularly interested in the sign of 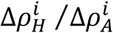 as this determines how an increased population of the *i*th conformation by the ligand-free protein 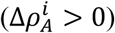 would lead to an increased 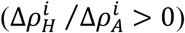 or decreased 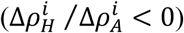 probability of forming the *i*th conformation with ligand. First, by dividing Eq. 31 by Eq. 30, we obtain

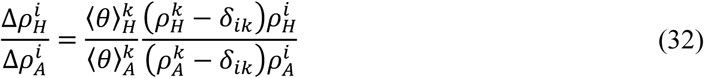

From Eq. 32 we see that 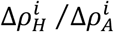 has the same sign for every conformation *i* and is solely determined by the sign of 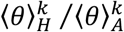. Note that the sign of 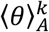 and 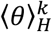 is determined by the integral in the numerator of Eq. 21 and 24, respectively.

When protein configurations that belong to the *k*th conformation (i.e., ***x*** ∈ Ω_*k*_) are structurally similar, it is reasonable to assume that when the intra-protein interactions are modified, the energy of each configuration of the *k*th conformation is shifted in the same direction. For instance, increasing the strength of an interaction that is present in the *k*th conformation, will likely lower the energy of all configurations in Ω_*k*_. Note that due to small structural differences between the configurations in Ω_*k*_, the energy of some of these configurations might be decreased slightly more than the others. Formally, shifting the energy of every ***x*** ∈ Ω_*k*_ in the same direction can be expressed as: either *θ*(***x***) ≥ 0 for every ***x*** ∈ Ω_*k*_ or *θ*(***x***) ≤ 0 for every ***x*** ∈ Ω_*k*_. Under these conditions, 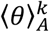 and 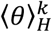 have the same sign (see Eq. 21 and 24), so from Eq. 32 it follows that for every conformation *i*

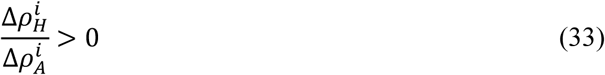

In conclusion, when the configurations of the *k*th conformation are either all increased or decreased in energy (and some might remain unaltered), then a shift in the ligand-free conformational equilibrium biases the ligand-bound conformational equilibrium in the same direction, i.e., 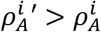 would lead to 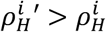 and 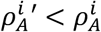 would lead to 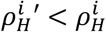 for every conformation *i*.

### 3.6 Constant protein-ligand interactions

Next, we will provide the result when (i) the energy of only the *k*th conformation is altered and (ii) the protein-ligand interaction energy is approximately constant for the *k*th conformation. When the protein-ligand interaction energy is approximately constant for the *k*th conformation, we have *g*(***x***) ≈ *g*_*k*_ for ***x*** ∈ Ω_*k*_, where *g*_*k*_ is a constant. Because of the structural similarity between the different configurations of a conformation, it seems reasonable that only small differences exist in the protein-ligand interaction energies for every ***x*** ∈ Ω_*k*_, and thus be approximately constant. When this is the case, we have

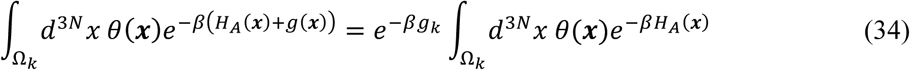

and

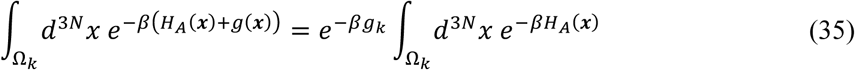

By dividing Eq. 34 by Eq. 35 and using Eq. 21 and 24 we obtain

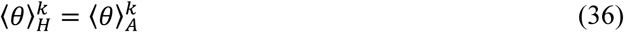

Note that in the simplest case when *θ*(***x***) is a constant for ***x*** ∈ Ω_*k*_, as in the example of Section 3.3, but without making any assumption about *g*(***x***), Eq. 36 also holds. In both cases, Eq. 32 becomes

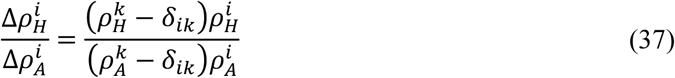

Since the right-hand side of Eq. 37 is strictly positive for every *i* (assuming 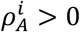 for every *i* ∈ {1, …, *m*}) it follows that 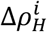 is positively related to 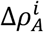. Thus, 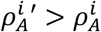 would lead to 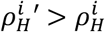 and 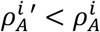 would lead to 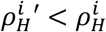 for every conformation *i*.

Up to this point, we ignored the fact that proteins in their physiological environment are surrounded by a large number of solvent molecules. Previous studies showed that the solvent degrees of freedom can be simply integrated out and that this results into the addition of a solvation free energy term to the energy of each configuration^35, 50^. In our case, this means replacing *H*_*A*_(***x***) with *H*_*A*_(***x***) + *w*(***x***) and replacing *H*_*A*_(***x***) + *g*(***x***) with *H*_*A*_(***x***) + *g*(***x***) + *v*(***x***), where *w*(***x***) and *v*(***x***) are the solvation free energy terms of the ligand-free and ligand-bound protein configuration, respectively. The addition of a solvation free energy term does not affect any of our conclusions, except the condition for which Eq. 36 is true. In the presence of solvent, not only does *g*(***x***) need to be approximately constant over Ω_*k*_, but also *w*(***x***) and *v*(***x***), in order to obtain Eq. 36. We argued above that a constant *g*(***x***) over Ω_*k*_ is expected to arise from the structural similarities of the configurations in Ω_*k*_. It seems logic that these structural similarities would then also give rise to similar protein-solvent interactions and thus lead to approximately constant *w*(***x***) and *v*(***x***) terms over Ω_*k*_.

### 3.7 Fraction of proteins having a ligand bound

So far, our analysis focused on the conformational equilibrium of a ligand-free and a ligand-bound protein. Here, we analyse how the fraction of proteins occupied by a ligand is affected by changes in the energy of a single conformation. To do this, we couple the protein to a reservoir which contains *L* ligand molecules within a volume *V*^51^. Within the grand canonical ensemble, the protein can exchange ligands and energy with the reservoir so that it can acquire all the *m* conformations with and without ligand^34^. The grand partition function of the protein is

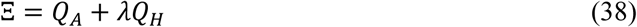

where *λ* = *e*^−*βμ*^ is the fugacity and *μ* the chemical potential. The fugacity is a function of *L*, with *λ* → 0 when *L* → 0 and *λ* → ∞ when *L* → ∞.

In the presence of *L* ligands, the equilibrium probability that the protein has a ligand bound is

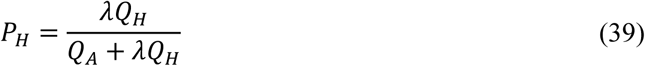

and the probability that the protein is ligand-free is then simply *P*_*A*_ = 1 − *P*_*H*_.

If we assume the ligand solution behaves as an ideal solution, i.e., the ligand molecules are non-interacting particles, then *λ* = *λ*_0_*L*, where *λ*_0_ is the standard fugacity. Under these conditions, *P*_*H*_ reduces to the Hill-Langmuir equation^52^, i.e.,

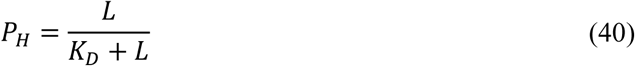

where the dissociated constant *K*_*D*_ (the binding or association constant is 1/*K*_*D*_) is equal to

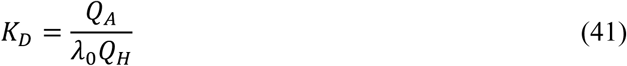

By using Eq. 2 and 5 this can be expressed as

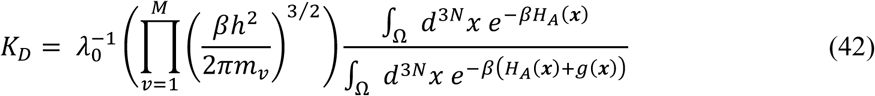

where *m*_*v*_ is the mass of the *v*th atom of the ligand. When *L* ≫ *K*_*D*_ most proteins have a ligand bound, when *L* ≪ *K*_*D*_ most are ligand-free and when *L* = *K*_*D*_ half of the proteins have a ligand bound. Therefore, *K*_*D*_ determines the affinity between the protein and the ligand.

When the Hamiltonian is changed by a perturbation *ϵθ*(***x***), the new *K*_*D*_ becomes

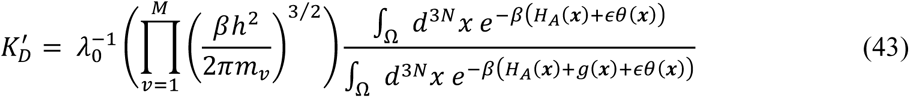

and the new probability that a protein has a ligand bound is

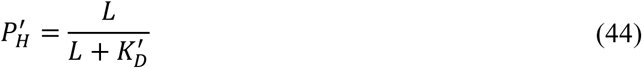

To first-order in *ϵ* we have

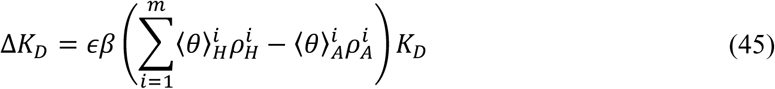

where 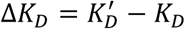. When only the *k*th conformation is altered (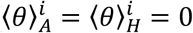 for every *i* ≠ *k*) this becomes

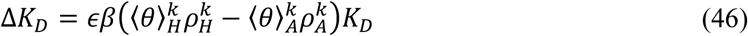

In case of the example of Section 3.3, we have

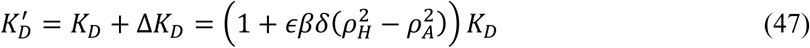

where, in this example, 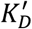 and *K*_*D*_ are the dissociation constants of the mutant and wild type protein, respectively. In the equilibria as depicted in Figure 2 we have that 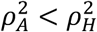, so when *δ* < 0 the affinity for the ligand increases 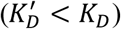 by the mutation and when *δ* > 0 we have 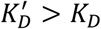. If *δ* < 0 we concluded from Eq. 26 that 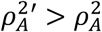 and when *δ* > 0 that 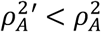. In other words, if the mutation shifts the ligand-free conformational equilibrium towards conformation 2, the affinity for the ligand increases and it decreases when the equilibrium is shifted towards conformation 1.

The above example suggest that a relationship between Δ*K*_*D*_ and 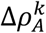 exists. To investigate this further, we can combine Eq. 30 and 46,

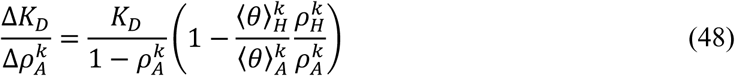

Now if

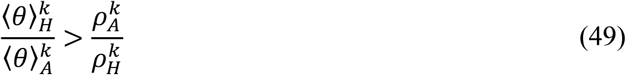

then

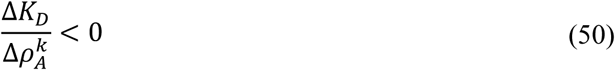

Thus, when Eq. 49 holds, Δ*K*_*D*_ and 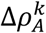 are negatively related. In other words, when 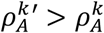, then the affinity between the protein and the ligand would increase 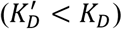 and when 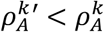 the affinity would decrease 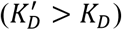. The condition that either *θ*(***x***) ≥ 0 for every ***x*** ∈ Ω_*k*_ or *θ*(***x***) ≤ 0 for every ***x*** ∈ Ω_*k*_, ensures that 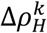 and 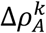 are positively related (Section 3.5), but it does not imply that Δ*K*_*D*_ and 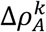 are negatively related. However, when the protein-ligand interactions are approximately constant for the *k*th conformation and/or *θ*(***x***) is a constant, we have 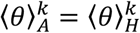 (Section 3.6), so that Eq. 48 becomes

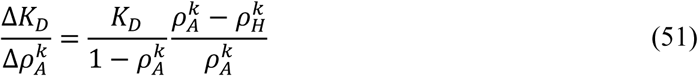

From Eq. 51 it follows that Δ*K*_*D*_ and 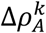 are positively related when 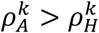 and are negatively related when 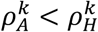. These results show that changes in the ligand-free conformational equilibrium can increase or decrease the affinity for the ligand and that this depends on the conformational equilibria of the ligand-free and ligand-bound protein.

## 4 DISCUSSION

By using classical statistical mechanics and assuming that the Hamiltonian function is additive, we showed how changes in the ligand-free conformational equilibrium affect the ligand-bound conformational equilibrium. We find that, under certain conditions as stated in the result section, a shift in the ligand-free conformational equilibrium biases the conformational equilibrium of the ligand-bound protein in the same direction. Furthermore, changes in the ligand-free conformational equilibrium can increase or decrease the affinity for the ligand. Our analysis is concerned only with equilibrium quantities (free energies, average structure etc.) and is thus independent of any kinetic pathway that connects the conformational states. Therefore, our analysis is valid irrespective of the ligand-binding mechanism, i.e., if the ligand binds via the induced-fit^5^, conformational selection^25^ or any other mechanism^53^. Furthermore, the results do not only cover the case in which a protein binds a simple ligand molecule, but would also apply to enzyme-substrate, protein-DNA, protein-RNA, protein-protein interactions and our results also apply to soluble proteins as well as membrane proteins.

From an intuitive perspective, when the stability of a ligand-free conformation is altered while at the same time leaving the protein-ligand interactions unchanged, then the stability of that conformation with a ligand bound should be shifted in the same direction. Indeed, a strict positive relation between 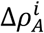 and 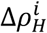 exists when the energy of only a single protein configuration ***x*** is altered (the numerator of Eq. 21 and 24 have the same sign when *θ*(***x***) is a Dirac delta function). A protein conformation consists of an enormous number of configurations that are structurally highly similar. So altering the energy of one configuration will most definitely also affect the others. Therefore, we considered changes in the energy of a set of configurations that all belong to the same protein conformation (i.e., the *k*th conformation). We showed that when the energies of all the configurations of the *k*th conformation are shifted in the same direction, i.e., they are either all increased or all decreased in energy (although some might remain unchanged), then a positive relation between 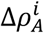 and 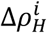 is predicted to exists (Section 3.5). The assumption that the energies are shifted in a similar direction seems reasonable from a structural point of view (see also Section 3.5). The structure of the configurations of a protein conformation are expected to be highly similar. Therefore, increasing the strength of an interaction is expected to lower the energy of all the configurations that belong to a conformation, rather than increasing the energy of some of these configurations and decreasing the energy of the others. The positive relation between 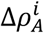 and 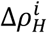 could be tested experimentally, by examining how changes in the ligand-free and ligand-bound conformational equilibrium are related when the intra-protein interactions, which are uniquely formed in a particular conformation, are altered, for example, by a point mutation.

When only the energy of the *k*th conformation is altered and, in addition, the protein-ligand interaction energy is approximately constant for the *k*th conformation, a simple relationship between the conformational equilibria is predicated to exist. First, 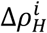 is positively related to 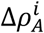, so increasing 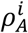 will increase 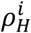 and vice versa (Section 3.6). Secondly, when the intra-protein interactions of the *k*th conformation are altered such that 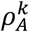 increases, then *K*_*D*_ will decrease when 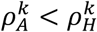 and increase when 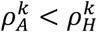 (Section 3.7). Alternatively, when 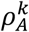 decreases, then *K*_*D*_ will decrease when 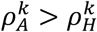 and increase when 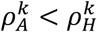. Of course, not only the ligand-free conformational equilibrium, but also the direct protein-ligand interaction energy *V*(***x***′, ***x***) determines the *K*_*D*_ and the ligand-bound conformational equilibrium. However, proteins with a more pronounced sampling of a particular ligand-free conformation, would require less strong protein-ligand interactions to form this conformation with ligand. This shows how a protein might fine-tune its function by balancing the energetic properties of its structure and the interactions with the ligand.

The predictions of this work are only qualitative, however, we believe that they are experimentally testable and already a number of observations seem to support them. Examples include the SBPs, which form a large class of structurally related proteins that are associated with ABC importers^54^, tripartite ATP-Independent periplasmic (TRAP)^55^ transporters and other systems^56, 57^. These proteins consist of two subdomains connected by a flexible hinge. Ligand binds between the two subdomains, causing them to come closer together and shift the conformational equilibrium from an open to a closed conformation^10, 11^. Seo et al. designed various point mutations to destabilize the open conformation of the SBP MalE^58^. They found a positive correlation between the affinity for the ligand maltose and the sampling of the closed conformation by the ligand-free protein. This is consistent with the prediction of Eq. 51. First, destabilising the open conformation would result in 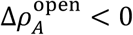, as was observed in their work^13, 58^. Secondly, in all their mutants it was observed that 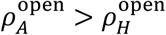, so when the protein-ligand interactions are approximately constant for the configurations of the open conformation, then we predict that Δ*K*_*D*_ < 0 since 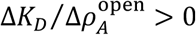, which is in agreement with their findings^58^. Marinelli et al. studied the SBP TeaA using computational methods and X-ray crystallography^59^, and showed that in the absence of the ligand ectoine the closed conformation is ∼5 *k*_*b*_*T* higher in energy than the open conformation. By making a triple alanine mutant to destabilize the closed conformation, the ligand-free closed conformation becomes now ∼24 *k*_*b*_*T* higher in energy than the open conformation. Furthermore, the mutations increased the *K*_*D*_ 50-fold, from 200 nM to 10 μM. This is again consistent with the predictions made, i.e., destabilizing the closed conformation leads to 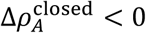 and since 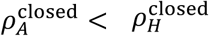 it follows from Eq. 51 that 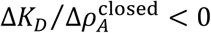.

Another example is the *Escherischia coli* enzyme Adenylate Kinase (AK) that catalyses the conversion of ATP and AMP into two molecules of ADP^60^. AK populates two conformations, an open and closed conformation^61^. Closing is required for the chemical reaction, while opening allows ATP and AMP to bind and ADP to be released. Structural analysis with NMR and smFRET demonstrated that the nucleotide-free conformational equilibrium of AK lies towards the open conformation^18, 62^. By breaking a hydrogen bond that is uniquely formed in the closed conformation, the conformational equilibrium of nucleotide-free AK is further biased towards the open conformation. Furthermore, the mutation decreased the affinity of AK for nucleotides. This is consistent with Eq. 51, since we have that 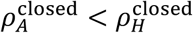, so 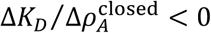 and thus Δ*K*_*D*_ > 0, since 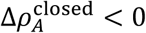.

A final example is the soluble ABC protein ABCE1, that facilitates ribosome recycling in archaea and eukaryotes, by splitting stalled ribosomes into its large and small subunits^63^. In our previous work we showed that ABCE1 is in an equilibrium between three conformations: open, intermediate and closed^12^. Interactions with ligands such as nucleotides and ribosomal subunits influence the conformational equilibrium. In our previous work we showed that ABCE1 derivatives with a N-terminal truncation have the conformational equilibrium shifted towards the intermediate conformation under all conditions (free, nucleotide, ribosome and nucleotide/ribosome)^12^. This is consistent with our prediction, namely that changes in 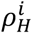 are positively related to changes in 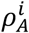.

Taken together, the results in this work provide testable hypotheses how the ligand-free and ligand-bound conformational equilibrium are related and how changes in these equilibria affect the affinity of the protein for the ligand.

## 5 ACKNOWLEDGEMENTS

This work was financed by an ERC Starting Grant (No. 638536 – SM-IMPORT to Thorben Cordes) and done during work in the Cordes lab at the University of Groningen. I thank Thorben Cordes, Giorgos Gouridis, Monique Wierstsma and Erik van der Giessen for fruitful discussions and valuable input.

